# Single-cell proteomics: a powerful new tool to study kidney cell heterogeneity

**DOI:** 10.1101/2025.09.23.678003

**Authors:** Qi Wu, Lei Cheng, Adrienne M. Assmus, Xiang Zheng, Cristina Esteva Font, Charlotte C. Petersen, Xiangning Ding, Søren R. Paludan, Robert A. Fenton

**Affiliations:** Department of Biomedicine, Aarhus University, Aarhus, Denmark

**Keywords:** single-cell proteomics, kidney, distal convoluted tubule, cell heterogeneity, NCC, *Slc12a3*

## Abstract

**Background:** Technological advancements in protein mass spectrometry have significantly enhanced analytical sensitivity and throughput, enabling single-cell proteomics by mass spectrometry (SCP-MS) to become reality. SCP-MS allows high-resolution analysis of cellular heterogeneity and function, bypassing bulk analysis limitations. Here, we used SCP-MS to document at the protein level the cellular diversity of mouse kidney cells. We further focused SCP-MS on low abundance distal convoluted tubule (DCT) cells that are essential for body electrolyte homeostasis and blood pressure control.

**Methods:** Mouse kidney cells were isolated via enzymic digestion and single-cells isolated using fluorescence-activated cell sorting (FACS). DCT cells were isolated from mice with GFP expression specifically in the DCT (parvalbumin-GFP) using a similar workflow. SCP-MS analyses was performed using an Orbitrap Ascend Mass Spectrometer with a FAIMS Pro Duo interface coupled with Wide ISolation window High-resolution MS1 Data Independent Acquisition (WISH-DIA). Spectronaut was used for protein identification and quantification, and cell-type annotation and clustering were performed in Python by scanpy implementation.

**Results:** Workflow benchmarking using Hela cells confirmed a successful SCP-MS setup. From 768 single mouse kidney cells, SCP-MS identified 2626 unique proteins. Computational approaches resolved various nephron segments, including distinct DCT populations. From enriched DCT cells, 1912 proteins were identified, enabling classification of three populations - DCT1, DCT2, and a proliferative subset (ProLIF) that represented a transient state between DCT1 and DCT2. ProLIF cells had elevated abundance of the sodium-chloride cotransporter NCC and represented approximately 33% of DCT cells, substantially exceeding previous transcriptomic estimates of ProLIF cell frequency (~0.1%). High proliferation (~39%) of DCT cells was confirmed using immunohistochemistry.

**Conclusions:** This SCP-MS analysis of mouse kidney uncovered significant cellular heterogeneity not captured effectively using transcriptomics. Despite imitations in proteome depth and throughput, SCP-MS provides a powerful approach for investigating kidney cellular dynamics at the protein level.

**Key points:**

- We developed a single-cell proteomics by mass spectrometry (SCP-MS) workflow identifying up to ~2,000 proteins in a single mouse kidney cell
- SCP-MS identified a substantial proliferative subset of cells in the DCT underestimated by transcriptomics
- SCP-MS could inform on clinical strategies for kidney disease monitoring, offering robust indicators of tubular injury or regeneration.

## Introduction

Single-cell proteomics by mass spectrometry (SCP-MS) has emerged as a transformative tool for unraveling cellular complexity at unprecedented resolution ^1,2^. Unlike bulk proteomics, SCP-MS analyzes individual cells, providing a deeper understanding of cellular heterogeneity, state-specific functionalities, and dynamic biological processes.

To achieve SCP-MS, several technological advancements have occurred in recent years that have significantly expanded the proteomic coverage, throughput, and quantitative precision achievable at the single-cell level. One such advancement is in commercial mass spectrometers, with several SCP-capable instruments now on the market ^3^. These state-of-the-art instruments perform data-independent acquisition (DIA) ^4,5^ and have allowed researchers to employ label-free approaches using novel chromatographic columns and short chromatography times to enhance throughput ^5^. High-field asymmetric waveform ion mobility spectrometry (FAIMS) ^6^ can further increase sensitivity on some instruments. The use of high-resolution-MS1 coupled with wide-window DIA, termed WISH-DIA (Wide Isolation window High-resolution MS1-DIA) ^4^, has also proved advantageous for SCP-MS due to its capacity to accommodate long ion injection time and high resolution under moderate scan speeds.

The kidney maintains water, electrolyte and acid-base homeostasis, helps regulate blood pressure, removes waste products and secretes hormones. To perform all these functions, the kidney has a complex cellular composition of at least 16 different cell types ^7-9^, including podocytes, mesangial cells, macula densa cells, immune cells, interstitial cells and numerous endothelial cell subtypes. Furthermore, the nephron is comprised of highly specialized epithelial cell types organized into distinct renal tubule segments with unique proteomic landscapes ^10^. The distal convoluted tubule (DCT) epitomizes such a nephron segment of considerable functional importance in respect to renal electrolyte handling, in particular sodium and calcium reabsorption ^11^. However, due to its relatively low cellular abundance, proteomic assessment of the DCT has been limited ^12-14^, and hence cellular heterogeneity of this segment has until now been classified based on transcriptomic approaches ^15^.

The kidney represents an ideal organ system to leverage the strengths of SCP-MS, in particular it has the potential to significantly enhance our understanding of the heterogeneity of DCT cells. Therefore, here we generated a SCP-MS WISH-DIA workflow based on a Vanquish Neo HPLC coupled to a FAIMS Pro Duo-linked Orbitrap Ascend Tribrid mass spectrometer to perform SCP-MS on mouse kidney. After initial benchmarking of performance using commercial Hela cell digests or FACS sorted Hela cells, the workflow was applied to randomly isolated single cells from mouse kidney to generate for the first time single cell maps at the protein level. We subsequently performed SCP-MS on a DCT-enriched single-cell population, enabling classification of three populations - DCT1, DCT2, and a highly abundant proliferative subtype (ProLIF). Our studies demonstrate the power of SCP-MS for investigating kidney cellular dynamics at the protein level and capturing novel states and heterogeneity that may be missed using transcriptomics.

## Methods

A schematic overview of the entire SCP-MS workflow is depicted in ***Figure 1***.

**Figure 1.**
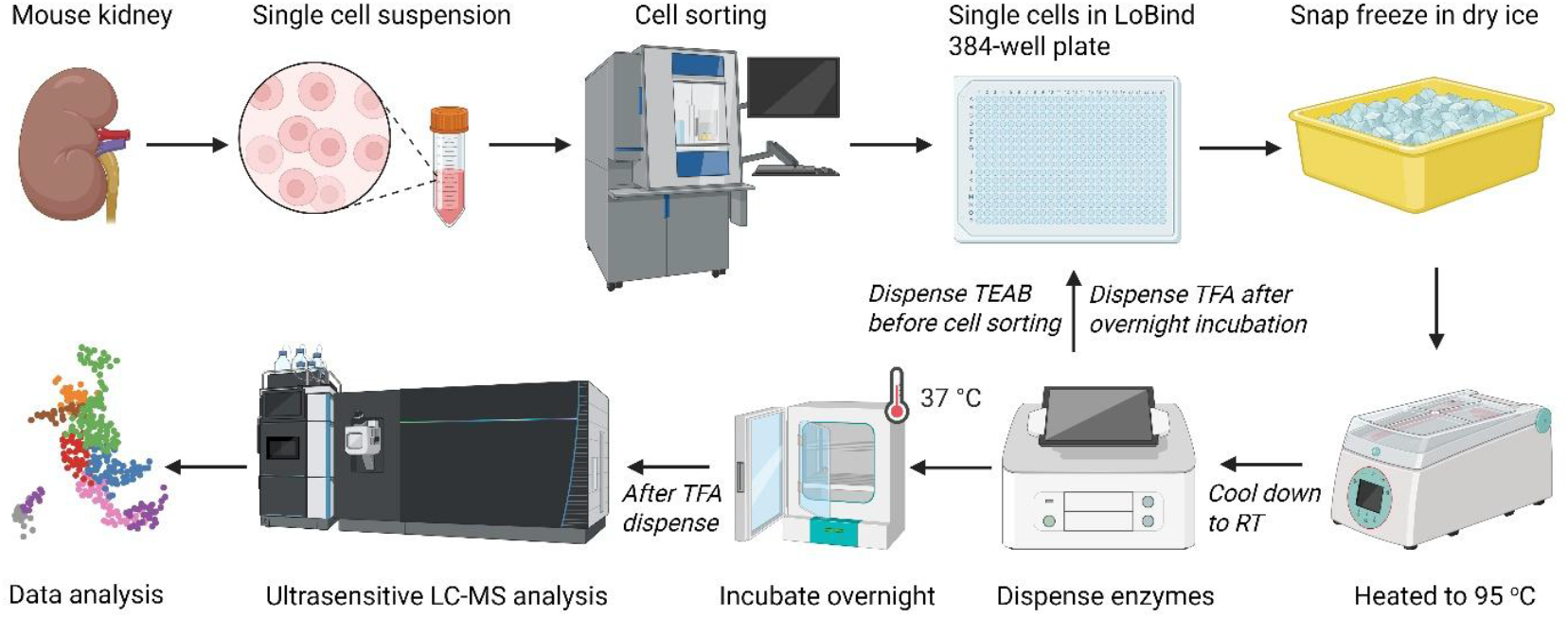
Schematic overview of the SCP-MS workflow. Fresh mouse kidney tissue is enzymically dissociated to a single-cell suspension, and individual cells are FACS-sorted into LoBind 384-well plates containing triethylammonium bicarbonate buffer. Cells are snap-frozen on dry ice, lysed at 95 °C for 5 minutes and then cooled to room temperature (RT). Proteolytic enzymes are dispensed to each well and digestion proceeds overnight at 37 °C. After reaction quenching with trifluoroacetic acid samples are analyzed with ultrasensitive nanoLC–MS analysis. The resulting proteome data are processed for cell-type assignment and downstream analyses.

### Hela cell culture

HeLa cells (ATCC CCL-2) were maintained at 37 °C in a humidified 5% CO_2_ incubator. Cells were cultured in Dulbecco’s Modified Eagle Medium (DMEM, high glucose, 4.5 g/L) supplemented with 10% (v/v) fetal bovine serum (FBS) and 1% penicillin–streptomycin. Media was replaced every 2–3 days and cultures were passaged at 70–90% confluence using trypsin–EDTA. For experiments, cells were at ~passage 20 and 60–80% confluent at the time of use. Cells were trypsinized, number and viability were determined by trypan blue exclusion (target viability ≥90%) and cells were stored in PBS immediately before sorting.

### Mice and study approval

Studies were conducted in compliance with Danish regulations, under an experimental animal license approved by the Animal Experiments Inspectorate, Ministry of Food, Agriculture, and Fisheries, Danish Veterinary and Food Administration (Dyreforsøgstilsynet, 2024-15-0201-01672). C57Bl/6JBomTac mice (referred to as C57Bl/6) were acquired from Taconic (Taconic Biosciences, NY). Transgenic mice expressing enhanced green fluorescent protein (eGFP) throughout the DCT ^16,17^ under the control of the parvalbumin (PvAlb) promoter (B6;129P2-Pvalbtm1(cre)Arbr/J) were kindly provided by Dr. H.Monyer (University of Heidelberg, Germany). The mice used have been backcrossed for 6 generations to C57Bl/6JBomTac mice. PvAlb-GFP mouse genotype was verified by PCR on genomic DNA from tail biopsies (primers: 5’-TAACTATGCGGCATCAGAGC-3’ and 5’-GCCTCCAGGTCGACTCTAGAG-3’) using standard techniques. All animals were kept in standard cages until use, in a room with a 12:12 h artificial light-dark cycle, a temperature of 21 ± 2ºC, and humidity of 55% ± 2%, with free access to water and standard rodent diet (1324 pellets, Altromin).

### Mouse kidney single cell suspension

Two male C57Bl/6 mice of 10 weeks old, and two male PvAlb-GFP littermates of 12 weeks old were used. Mice were anesthetized with isoflurane and perfused through the heart left ventricle with 10 ml of collagenase solution containing 2 mg/mL collagenase B (Roche COLLB-RO), 250 μg/mL collagenase IV (Gibco 17104-019), 125 mM NaCl, 0.4 mM KH_2_PO_4_, 1.6 mM K_2_HPO_4_, 1 mM MgSO_4_, 10 mM Na-acetate, 1 mM α-ketogluterate, 1.3 mM Ca-gluconate, 5 mM glycine, 30 mM glucose and 5 μg/mL DNase I (Sigma), pH 7.4. Mice were euthanized by cervical dislocation, the kidneys were rapidly excised, cut into small pieces of ~1mm^3^, and single cell suspensions prepared as previously described [17]. Briefly, the ~1 mm^3^ pieces were incubated at 37°C in more collagenase solution (described above) with continuous mixing at 850 rpm. After 10 min, half of the collagenase solution was removed and replaced with an equal volume of fresh solution, and samples were incubated similarly for a further 10 min. The samples were then centrifuged for 3 min at 200 *g*, and the supernatant removed. Cells were washed with a trypsin/EDTA solution (Lonza) containing 10 mM HEPES, 30 mM glucose and 50 µg/ml DNase I, resuspended in trypsin/EDTA solution and incubated for 15 min at 37°C with regular mixing. Cells were finally washed in DMEM/HamF12 cell culture medium (Gibco) containing 5% FBS, 30 mM glucose, 10 mM HEPES and 50 mg/ml DNase I before a final rinse and storage in PBS. Cell suspensions were immediately sorted at the FACS core facility, Aarhus University, Denmark.

### Cell sorting and in-situ sample preparation

Suspensions were passed through a 50 µm filter (BD Biosciences, Cat#340601) to eliminate cell clumps. To verify precise positioning of a single drop into a 384-well plate, an initial enzyme test was performed. A small fraction of the single cell suspension was resuspended in horse radish peroxidase (100 µg/mL, ChemCruz) and single cells from this suspension were sorted using a 100 µm nozzle and 20 psi on a 6-laser Bigfoot cell sorter with the SQS Software v.1.9.4 (Thermo Fisher) into a 384 well plate containing 2 µL 3,3’,5,5’-Tetramethylbenzidine (TMB) high sensitivity substrate (Biolegend). A substrate color change to blue verified accurate machine precision, allowing the cell suspension to proceed to sorting. Suspensions were propidium iodide stained (8 µg/ml, Invitrogen) to exclude dead cells and cells sorted using a single cell sort mask and straight down sorting, deflecting all events of no interest, directly into wells of a Lobind 384-well plate (Eppendorf skirted, 45 µL) containing 2 µL of 20 mM tetraethylammonium bromide (TEAB) buffer. For total kidney cells isolated from C57Bl/6 mice, cells were gated as non-debris, for singlets, and live cells. For cells from PvAlb-GFP mice, an additional gating for GFP fluorescence was included. Two examples of the FACS gating can be seen in ***Supplemental Figure 1***. Plates were centrifuged at 1,000*g* for 1 min, snap frozen on dry ice for 10 min and incubated for 5 min at 95°C in a heating block. A nanoliter level non-contact dispenser (iDOT, Cytena) was used to dispense 200 nL of Lys-C (10 ng/uL, Thermo Fisher, Cat#90307) to each well and the plate was covered and incubated at 37°C for 2 h. Subsequently, 200 nL of 10 ng/uL trypsin (Thermo Fisher, Cat#90057) was dispensed and the plate covered and incubated at 37°C overnight. Finally, 100 nL of 10% TFA was added to the wells, and after covering, the plate was stored at −80°C until LC-MS analysis.

### Ultrasensitive LC-MS analysis

Standard commercial Hela cell digest (Thermo Fisher, Cat#88329) or digested single cells in 384-well plates were analyzed on a Vanquish Neo HPLC coupled with Orbitrap Ascend MS and a FAIMS Pro Duo interface (Thermo Fisher). A 50 cm long low-load µPAC Neo column (Thermo Fisher) was used in a one-column setup. For Hela extract benchmarking and total kidney single cells, a 30-min gradient time was used for peptide separation with an initial flow rate ramping to 500 nL/min for 4 min, followed by one active separation window of 16 minutes of 5-22% of solution B (80% ACN and 0.1% FA) and another of 5 min of 22-40% of solution B at 200 nL/min, followed by a 5-min column rinsing phase with 99% solution B at 500 nL/min. MS analysis was done in DIA (Data Independent Acquisition) mode, with a FAIMS CV of −45. Parameters for MS1 scans include a scan rage of 375-1010, Orbitrap resolution of 240 K, maximum ion injection time of 507 ms, normalized AGC of 250%. Parameters for MS2 scans include DIA window size of 68 Da, Orbitrap resolution of 240 K, maximum ion injection time of 507 ms, normalized AGC of 1000%, collision energy of 32%, number of scan events = 9, and loop count of 3 (one MS1 scan inserted between every 3 MS2 scans). For single DCT cells, a slightly modified method was used. Specifically, a 25-min gradient time was used for peptide separation with an initial flow rate ramping to 500 nL/min for 2.1 min, followed by one active separation window of 10.5 minutes of 5-22.5% of solution B and another of 7.5 min of 22.5-40% of solution B at 200 nL/min, and a 4.9-min column rinsing with 99% B at 500 nL/min. MS analysis was done in DIA mode, with a FAIMS CV of −50. Parameters for MS1 scans include a scan rage of 400-800, Orbitrap resolution of 240 K, maximum ion injection time of 507 ms and normalized AGC of 250%. Parameters for MS2 scans include a DIA window size of 68 Da, Orbitrap resolution of 240 K, maximum ion injection time of 507 ms, normalized AGC of 1000%, collision energy of 28%, number of scan event of 6, loop count of 3, and a loop time of 3 seconds.

### MS data analysis

Raw MS data was analyzed by direct DIA+ mode in Spectronaut (v18) against a human uniprot database for standard Hela digest and Hela cells (downloaded on Feb 3, 2022) or a mouse uniprot database for total kidney cells (downloaded on March 7, 2024) and for DCT cells (downloaded on Jan 6, 2025). Peptide and protein false discovery rates (FDR) were set at 1%. For standard Hela digest, default parameters were used except the quantification level was set to MS1. For all other searches, carbamidomethylation of cysteine was removed from the default panel. Identified proteins and the quantification matrix were exported to csv files for downstream data analysis.

### Cell type clustering and cell type annotation

Cell type clustering and cell type annotation were done by Python using scanpy implementation ^18^ as follows:

### Data processing and quality control

Raw single-cell expression values were imported from the Spectronaut csv file. Marker gene lists ^8,15^ (***Supplemental Table 1***), curated per cell type, were read from an Excel spreadsheet and uniformly transformed to ensure consistent labeling. The count matrix was transposed and stored in an AnnData object, with each feature (gene) assigned a cleaned, unique name. Routine data pre-filtering was performed, and cells with fewer than 100 detected genes and genes expressed in fewer than two cells were removed. Remaining counts were logarithmically transformed and scaled (capped at 10) to mitigate the influence of highly expressed genes.

### Dimensionality reduction, clustering, and annotation

Principal component analysis (PCA) was performed using the top 20 components, followed by computation of a nearest-neighbors graph (k = 10) and Leiden clustering at a resolution of 2 for total kidney cells or 3 for DCT cells. Each cluster was scored against the curated marker gene sets (***Supplemental Table 1***): for each cell type, the mean expression of its marker genes produced a “score,” and cells were provisionally labeled by the highest-scoring cell-type signature. Clusters with exceptionally low scores (bottom 1%) were designated “Unknown.” The final cell-type annotation was used to generate two-dimensional embeddings via UMAP. For total kidney single cell annotations, in addition to Leiden clustering, the following was further applied (*Penalized scoring to resolve shared markers* and *Threshold selection by grid search*) to reduce false positives from shared markers and resolve cell types better.

### Penalized scoring to resolve shared markers

Positive and negative marker sets per cell type were constructed:

Positive set: curated markers for that cell type present in the dataset.

Negative set: union of markers from competing cell types that share ≥1 positive marker with the focal type (excluding its own positives).

For each cell and cell type c, a penalized score was computed:

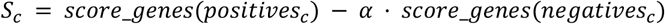

with α = 0.7. For each cell we recorded the best value and the margin between the best and second-best scores.

*Threshold selection by grid search*. Two decision thresholds were tuned on a grid: ABS_MIN (minimum acceptable best score; 0.10–0.85 in 0.05 steps) and MARGIN_MIN (minimum gap between top 2 scores; 0.05–0.45 in 0.05 steps). For each (ABS_MIN, MARGIN_MIN) pair, cells meeting both thresholds were reassigned from the baseline label to the new cell type. Each candidate was evaluated by a composite objective:

- Neighborhood Agreement (NAR): fraction of k-NN edges connecting cells with the same label after reassignment (higher is better).
- Mean confidence: mean best score among reassigned cells (higher is better).
- Label churn: fraction of cells whose label changed (lower is better).
- Unknown fraction: fraction of cells labeled “Unknown” (penalized only if “keep-unknown” is disabled).

The composite objective was:

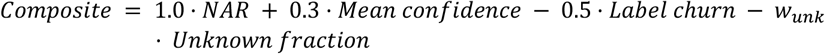

where w_unk_ = 0 when Unknown labels are kept (default), otherwise 0.2. The (ABS_MIN, MARGIN_MIN) pair maximizing this score was selected and applied dataset wide. The final label (cell_type_v2) equals the baseline cluster label except for cells that passed the tuned thresholds, which adopt their penalized score. Previous “Unknown” clusters remained “Unknown”.

### Differential expression and marker visualization

Within each Leiden cluster (for DCT SCP data) or re-annotated cell types (for total kidney SCP data), differentially expressed genes were identified by the Wilcoxon rank-sum test. All differential expression results were collated into a single table, sorted by scores. The first 10 unique proteins overall were used in the stacked violin plot for visualization.

### Trajectory inference

Diffusion pseudotime (DPT) analysis was initialized by selecting the first cell annotated as “DCT1” as the root. DPT values were computed, and pseudotime was visualized on UMAP coordinates with a continuous color scale. Additionally, PAGA (partition-based graph abstraction) was run on the cell-type graph to infer abstract connectivity, with both the PAGA graph and a UMAP overlay saved for interpretation. Scatter plots of selected genes (e.g. Mki67) against diffusion pseudotime were generated to assess expression trajectories. Boxplots (with jittered points) were used to display pseudotime distributions across predicted cell types.

### Immunofluorescence analysis

Formalin-fixed paraffin-embedded (FFPE) kidney tissue from an 8-week-old male C57BL/6JBomTac mouse (mouse #1) or two 14-week-old male C57BL/6JBomTac mice (mouse #2 and #3) were used for targeted validation. 5 µm sections were deparaffinized and rehydrated by sequential immersion three times in xylene, followed by a graded ethanol series from 99.6% to 70%. Antigen retrieval was performed by incubating the sections in 10 mM citrate buffer (pH 6.0) at 90 °C for 20 minutes. Sections were blocked in 5% bovine serum albumin (BSA) for 1 h at room temperature. Primary antibody incubation was carried out overnight at 4 °C using the following antibodies: Ki-67 (Santa Cruz, sc-23900, 1:100), PCNA (Sigma Aldrich, P8825, 1:20), or NCC (Stressmarc 402D, 1:1000). Sections were washed, incubated with 1:1000 of secondary antibodies (DAM647 and DAR555, Thermo Fisher) for 1 h at room temperature, counterstained with DAPI, and mounted using Thermo Fisher Diamond Antifade mounting medium. Imaging was performed using an Olympus VS120 microscope at 20× magnification, and images were saved in.vsi format. A non-primary antibody control was included as a negative control. Image analysis was conducted using QuPath. Whole-slide image annotations were created, and the ‘Positive Cell Detection’ plugin was applied with the following parameters: detection image set to fluorescence, requested pixel size of 0.5 µm, sigma of 1.5 µm, minimum area of 10.0 µm^2^, maximum area of 400.0 µm^2^, threshold of 100, maximum background of 1.0, split by shape set to true, include nuclei set to true, cell expansion of 3 µm, smooth boundaries set to true, and make measurements set to true. All other parameters were left at default values. The analysis was executed in batch mode, and results were exported in ‘detection’ format, allowing extraction of mean intensity values for subsequent quantification.

## Results

### Benchmarking of the LC-MS system and the sample preparation workflow

To assess the sensitivity of our LC-MS system for SCP-MS we performed benchmarking using 250 pg of Hela digest, which corresponds to the estimated total protein content of a single cell ^19^. A total of 2,100 protein groups (hereafter referred to as proteins) were identified (***Supplemental Figure 2A***) with excellent precision (***Supplemental Figure 2B***), showcasing the advantage of performing quantification on the MS1 level. Next, to examine the suitability of our workflow, we FACS sorted 100 single Hela cells into wells of a 384-well plate, performed our *in-situ* sample preparation and subjected them to LC-MS. On average, 1,756 proteins were identified (range of 720 to 2,150 proteins) (***Supplemental Figure 2A***). The numbers of proteins identified was in line with that seen by others using a similar workflow for SCP-MS ^4^.

### SCP analysis on total kidney cells

Once our workflows were validated, SCP-MS was performed on two full 384-well plates (n=768) of randomly sorted kidney cells. The total proteome reached a depth of 2,626 protein identifications (***Supplemental Table 2***), with an average of 899 proteins per cell and a range from 117 to 2,043 proteins. The data was clustered, annotated and visualized by Uniform Manifold Approximation and Projection (UMAP) in Python scripts through scanpy implementation ^18^ (see *Methods*). ***Figure 2A*** shows the cell-type annotations based on classic, well-established protein markers of specific cell classes in the kidney^8^. Ten distinct cell types, as well as a group labeled as “unknown” were identified. The relative expression of classical marker proteins can be seen in ***Figure 2B***, with some cell-type marker proteins e.g. nephrin (*Nphs1*) not identified, highlighting why podocytes were not annotated in UMAP. The top 10 markers with the highest differential expression scores, irrespective of cell type, were plotted in a stacked violin plot (***Figure 2C***). Apart from the protein markers SGLT2 (*Slc5a2)* and Megalin (*Lrp2*) used for annotation, we also identified Glycine amidinotransferase (*Gatm)*, Argininosuccinate synthetase (*Ass1*), Acyl-CoA Synthetase (*Acsm1*), Nudix Hydrolase 19 (*Nudt19*), and alanine aminotransaminase 1 (*Gpt1*) as secondary proximal tubule cell markers, and organic anion transporter 1 (*Slc22a6*) as a highly specific marker for proximal S2 cells. Aldo-keto reductase (*Akr1a1*) and Isocitrate dehydrogenase 1 (*Idh1*) showed relatively high expression across all cell types, with slightly higher expression in proximal tubule cells.

**Figure 2.**
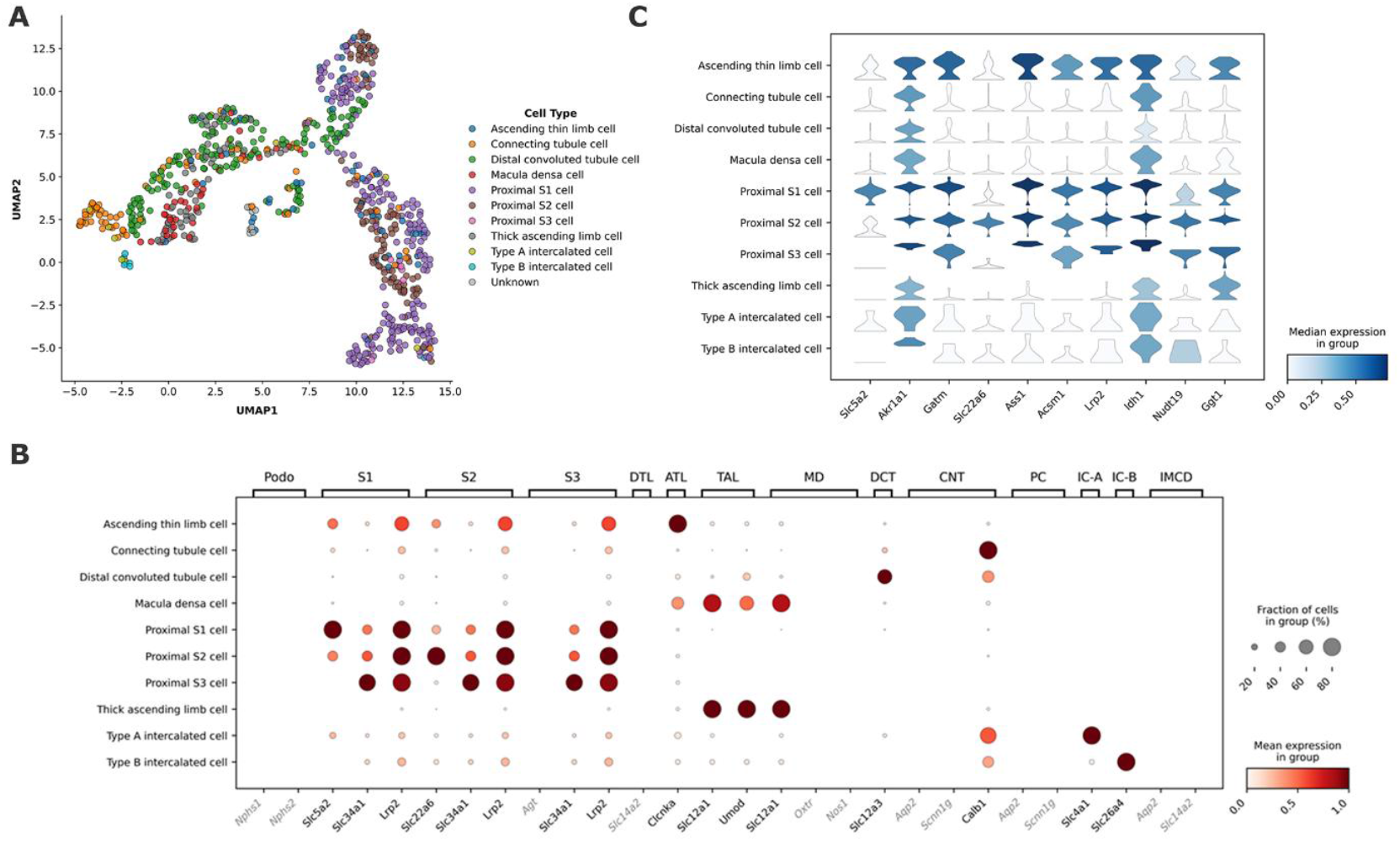
SCP-MS analysis of 768 randomly sorted cells from total kidney. (A) UMAP (Uniform Manifold Approximation and Projection) visualization of annotated cell types. (B) Expression of marker proteins. Upper x-axis are pre-defined cell types and lower x-axis are their marker proteins (listed as gene names). The y-axis is the annotated cell type. Greyed marker proteins (gene names) were not identified in the current SCP-MS data. Fill color encodes the mean expression in that group. (C) Stacked violin plots of the top 10 differentially expressed proteins across kidney cell types. For each mini-violin, width reflects the distribution of per-cell expression (density) and fill color encodes the median expression in that group.

### SCP analysis of distal convoluted tubule cells

Next, we performed SCP-MS analysis on 758 DCT cells isolated using FACS from the parvalbumin-GFP^+^ mouse. The purity of the sorted cells was 100%. A total of 1,912 proteins were identified (***Supplemental Table 3***), with an average of 846 proteins per cell and a range of 174 to 1,367 proteins. To elucidate cell-type heterogeneity and validate findings previously reported through single-cell transcriptomics, we used marker proteins from three distinct DCT populations [14] to annotate our DCT SCP-MS data: DCT1, DCT2, and a proliferative (ProLIF) subset (***Figure 3A***). Expression patterns of canonical markers separated these populations, with DCT1 characterized by high expression of NCC (*Slc12a3*), parvalbumin (*Pvalb*) and epithelial growth factor (*Egf*), whereas DCT2 cells had elevated levels of calbindinD28K (*Calb1*), calbindinD9K (*S100g*) and Kalikrien-1 (*Klk1*) (***Figure 3B***). The proliferative subset uniquely expressed proliferation-associated markers, notably DNA topoisomerase II (*Top2a*) and Ki-67 (*Mki67*). Interestingly, even though the sodium-chloride cotransporter NCC (*Slc12a3*) is a golden marker for DCT cells in general (***Figure 3C***), its expression was markedly higher in ProLIF cells compared to DCT1 or DCT2 subsets (***Figure 3B***). The top 10 markers with the highest differential expression scores in DCT subclasses included four markers used for annotation (*Mki67, S100g, Calb1*, and *Top2a*), but also secondary protein markers for ProLIF cells e.g. α and β propionyl-CoA carboxylase (*Pcca, Pccb)* and ATP-binding cassette subfamily A member 2 (*Abca2*) appears as a selective marker for DCT2 cells (***Figure 3D***).

**Figure 3.**
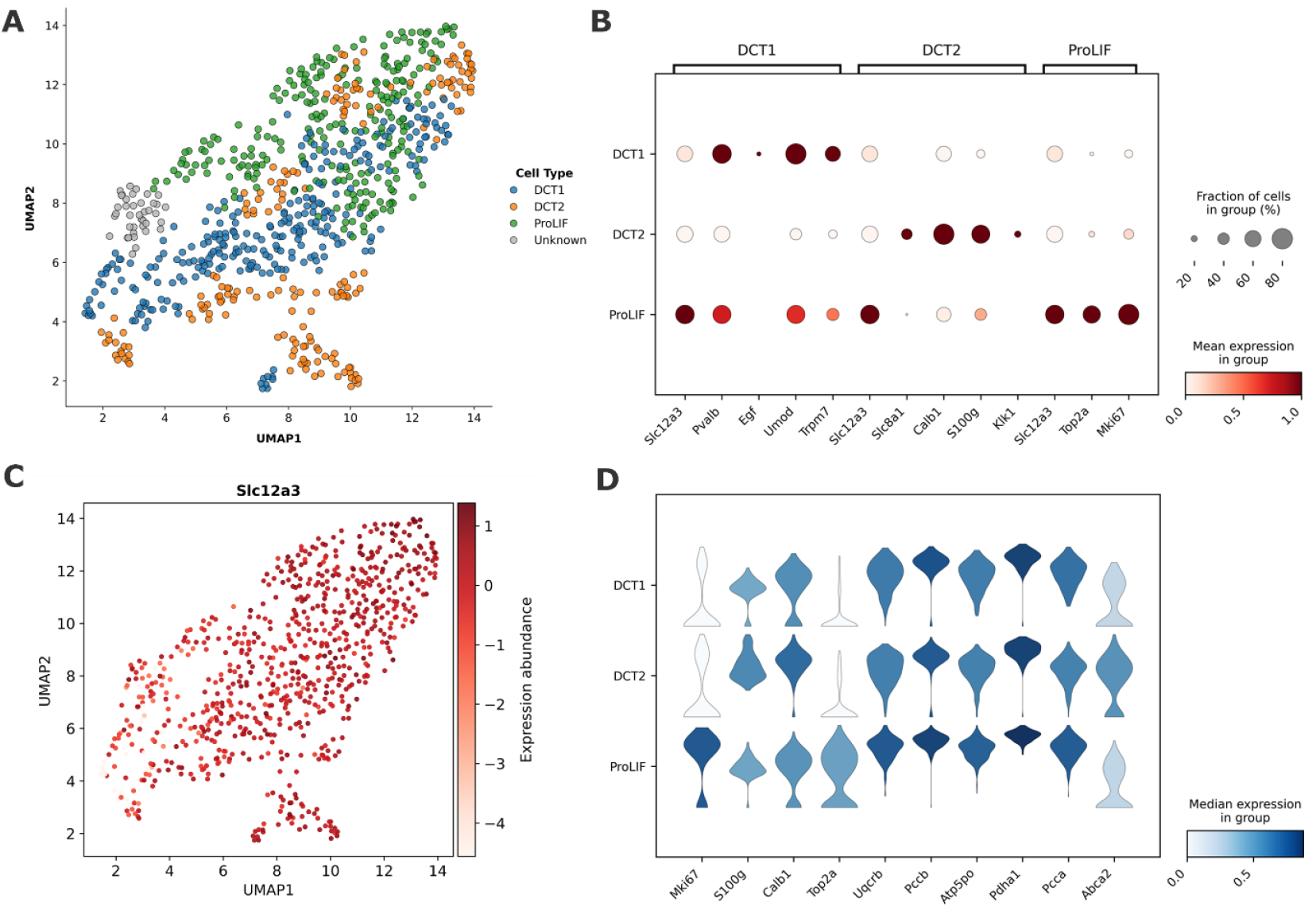
SCP-MS analysis of 758 DCT cells. (A) UMAP (Uniform Manifold Approximation and Projection) visualization of annotated DCT cell types. (B) Abundance of marker proteins in annotated cell types. Upper x-axis are pre-defined cell types and lower x-axis are their marker proteins (listed as gene names). The y-axis is the annotated cell type. Fill color encodes the mean expression in that group. (C) Golden DCT protein marker NCC (*Slc12a3*) abundance across all cells. (D) Stacked violin plots of the top 10 differentially expressed proteins across DCT cell types. For each mini-violin, width reflects the distribution of per-cell expression (density) and fill color encodes the median expression in that group.

Multiple trajectory analyses highlighted that the ProLIF cluster occupies an intermediate position between DCT1 and DCT2. When DCT1 is set as time zero, DCT1-derived cells span the earliest pseudotime values, DCT2 cells accumulate at the latest pseudotime values, and ProLIF cells appear squarely in the middle of this continuum (***Figure 4A***). Crucially, Ki-67 expression, which marks proliferative activity, peaks precisely in a similar mid-pseudotime window (***Figure 4B***). The partition-based graph abstraction (PAGA) graph places ProLIF cells topologically between DCT1 and DCT2 (***Figure 4C***), and the boxplots exhibit a narrow interquartile range of pseudotime between DCT1 and DCT2 (***Figure 4D***). Taken together, these patterns support a model in which ProLIF cells represent a transient, intermediate cell state along the differentiation continuum from DCT1 toward DCT2.

**Figure 4.**
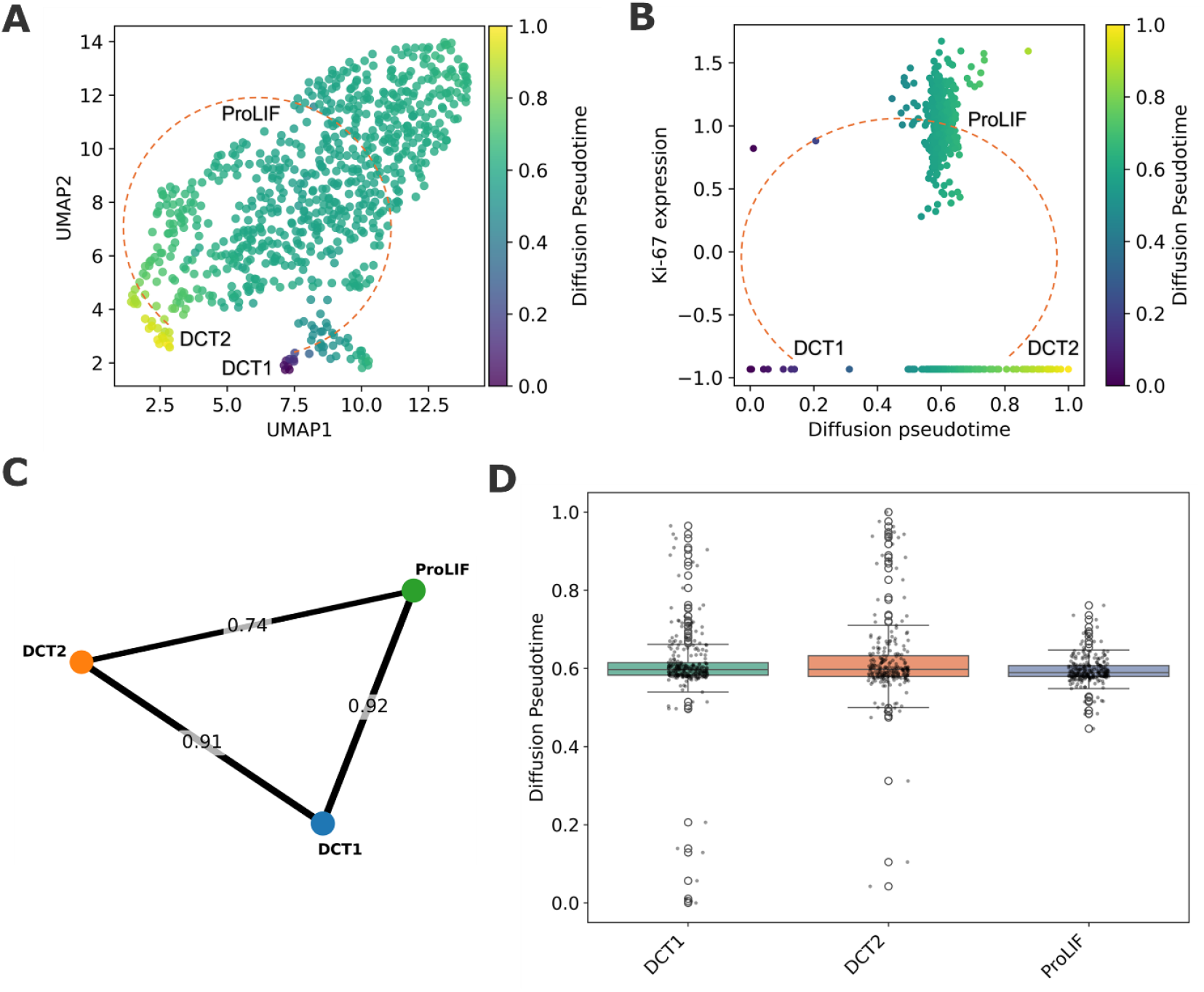
Integrated pseudotime and trajectory analyses of DCT1, DCT2 and ProLIF cell populations. (A) UMAP embedding of all cells, colored by diffusion pseudotime (rooted in DCT1), demonstrating a continuous progression from early (purple) to late (yellow) states. (B) Scatterplot of single-cell Ki-67 expression versus diffusion pseudotime. Ki-67 levels rise sharply in mid-pseudotime before declining, indicating maximal proliferative activity in the intermediate ProLIF state. (C) PAGA-derived abstract connectivity graph of the three clusters. Node size reflects cluster cell count; edge thickness (shown on the line) encodes connectivity strength, revealing strong pairwise links consistent with ProLIF as a transient intermediate. (D) Boxplots (with overlaid individual points) of diffusion pseudotime by annotated cell type. ProLIF cells exhibit a pseudotime between DCT1 and DCT2, with a narrow interquartile range, supporting their status as a transient cell type along the DCT1 towards DCT2 trajectory.

### Validation of Ki-67 in distal convoluted tubule

ProLIF cells comprised approximately 33% of the DCT cells identified in our SCP-MS dataset, which is significantly higher than the ~0.1% population of ProLIF cells reported by single-cell transcriptomics [14]. To validate our SCP data, immunofluorescence (IF) staining of Ki-67 in DCT cells across whole kidney tissue sections was performed. Semi-quantitative image analysis demonstrated that ~39% of DCT cells were Ki67-positive, consistent with the high proliferative activity suggested by our SCP-MS data (***Figure 5***). To further support this observation, we performed additional IF staining for the proliferation marker PCNA (***Supplementary Figure 3***). A similarly high number of DCT cells were PCNA positive, corroborating the Ki67-based cell quantification. These findings collectively strengthen the evidence for highly active proliferation in DCT cells, suggesting that the DCT may be constantly undergoing dynamic remodeling or regeneration even under homeostatic conditions.

**Figure 5.**
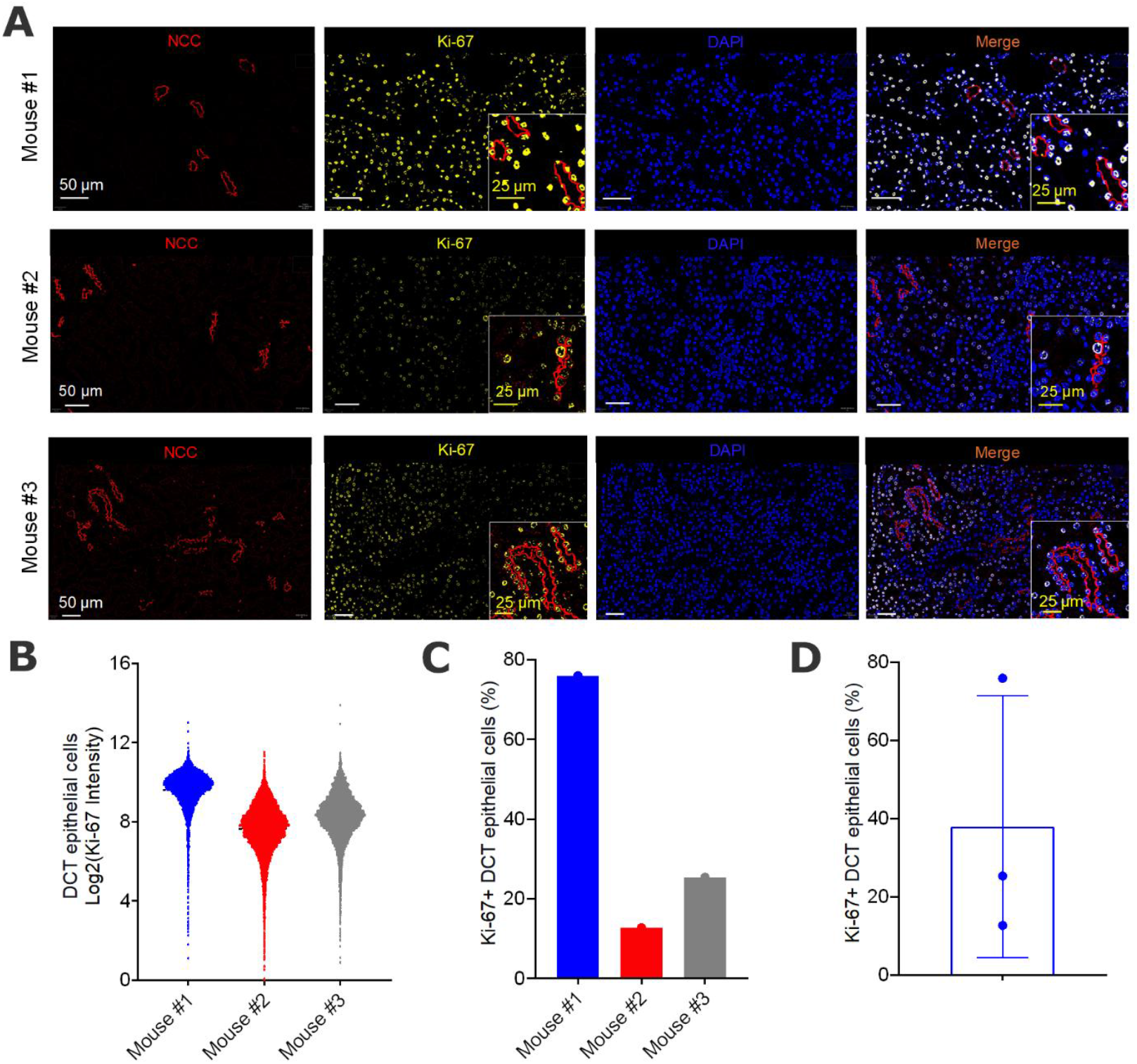
Immunofluorescence analysis of Ki-67 expression in distal convoluted tubule. (A) Representative images from three mouse kidneys of immunofluorescence staining of NCC (red) and Ki-67 (yellow) staining, alongside a nuclear DAPI counterstain (blue). (B) Staining intensities for Ki-67 within the NCC positive population (DCT) in three mouse kidneys. (C) Percentages of Ki-67 positive DCT cells compared to all DCT cells in each individual mouse kidney section. (D) Average data for Ki-67 positive DCT cells (mean ± S.D) across the three mice.

## Discussion

This study represents the first SCP-MS analysis of kidney, serving as a crucial proof-of-concept of the capability of SCP-MS to capture cellular heterogeneity within kidney cells on the protein level. This concept is highly evident from our data originating from DCT cells, where the proportion of ProLIF cells identified using SCP-MS was much higher compared to transcriptomics [14]. In addition, our results highlight how the SCP-MS approach may better capture transient cell states, given that proteins typically have longer half-lives compared to transcripts ^20^, potentially making proliferative cell states more detectable. Furthermore, as transient, transitioning and degenerative cell states are hallmarks and/or contributors to kidney injury ^9,21,22^, SCP-MS may be better suited to identify the maladaptive states associated with a decline in kidney function or signatures of epithelial cell repair.

This study also suggests that SCP-MS has the capacity to discover novel cell-type markers that are weak, inconsistent, or entirely absent at the transcript level, further supporting the idea that protein and mRNA are decoupled by translation, turnover, and localization ^20^. Supporting this, several of the proteins identified by SCP-MS as ProLIF markers, including those endcoded by *Uqcrb, Pccb, Atp5po, Pdha1*, and *Pcca*, did not stand out in single-nucleus RNA-seq studies ^15^. These mitochondria-related metabolic enzymes often show sharp protein-level specificity even when transcripts are sparse or lost to dropout. Likewise, ATP-binding cassette subfamily A member 2 (*Abca2*), a transporter protein, was identified as a DCT2 marker yet showed little DCT signal in snRNA-seq ^23^.

A central limitation of our study is proteome depth at the single-cell level. Unlike transcriptomics, SCP-MS is not amplifiable and currently it only identifies a subset of a cell’s proteome. This means that the ability to resolve cell identities is constrained to populations for which one or more marker proteins are both expressed and detectable under our acquisition and analysis settings. Subtypes lacking robust proteotypic peptides (e.g. low-copy proteins, multi-pass membrane proteins, or markers with unfavorable peptide chemistry) are under-represented, and closely related states may collapse into broader classes. Therefore, it is necessary to integrate multiple annotation strategies to achieve comprehensive and accurate cell type assignments in SCP-MS studies. Importantly, non-detection of a specific protein should not be interpreted as true absence of expression; rather, it reflects sampling and identification limits inherent to SCP-MS. This limitation most likely explains the identification of only 10 different kidney cell types in this study, whereas transcriptomics suggests additional cellular diversity [20]. Another constraint is the low throughput of our current SCP-MS workflow, limited to approximately 32 cells per day, which poses a considerable time limitation compared to the extremely high number of cells measurable with single-cell transcriptomics. However, although SCP-MS will never have the capacity to profile several hundred cells per day, technological advances in instrumentation (e.g. Orbitrap Astral) are rapidly accelerating SCP-MS, enabling markedly deeper proteome coverage (range of 5 to 6,000 proteins) at much higher speed and throughput ^3,24^.

In perspective, integrating SCP-MS with single-cell RNA-seq approaches leverages the complementary strengths of the two modalities; transcriptome breadth and protein-level functional readouts, which can lead to more accurate cell annotations and mechanistic insights ^25^. In the kidney, such integration could result in sharpened nephron taxonomy, link transporter and metabolic protein abundances to mRNA-expression patterns, clarify mechanisms underlying electrolyte handling, or identify novel drivers of epithelial cell repair during kidney function decline. Furthermore, the rapid advancement of SCP-MS towards deeper coverage and higher throughput will provide a more complete and actionable view of kidney biology, accelerating progress in precision nephrology.

## Data availability

All MS data within this manuscript can be accessed at https://massive.ucsd.edu/ProteoSAFe/dataset.jsp?task=802717074a0946deb84a751c844d4fb5, with reviewer account: MSV000098593_reviewer, and password: eh6qA1fy426SOvB4. Python code for data analysis and associated data can be accessed at: https://github.com/qiwu-au/SCP_mouse_kidney.

## Supplementary Information

Supplementary material is available separately.

## Acknowledgements

The mass spectrometry equipment utilized in this study is part of the Danish Single-Cell Examination Platform (CellX) established with support from the Danish Research Agency Infrastructure Program (5229-0009B). Funding to R.A. Fenton is provided by the Novo Nordisk Foundation, the Lundbeck Foundation, and the Danish Medical Research Council. We thank Sune Jonathan Keidser-Nilsson and colleagues from the FACS Core Facility, Aarhus University, Denmark for assistance with FACS experiments. We would like to acknowledge the technical assistance of Inger Merete S. Paulsen and Tina Drejer.

