## Supplemental Information for "Single-cell proteomics: a powerful new tool to study kidney cell heterogeneity"

**A**

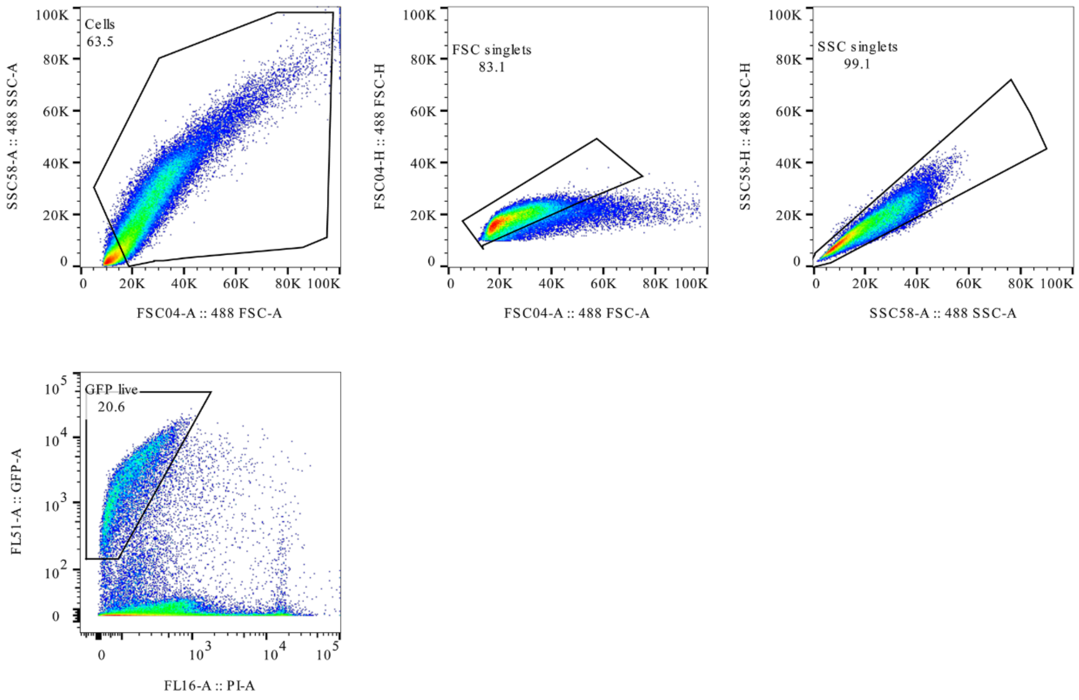

**B**

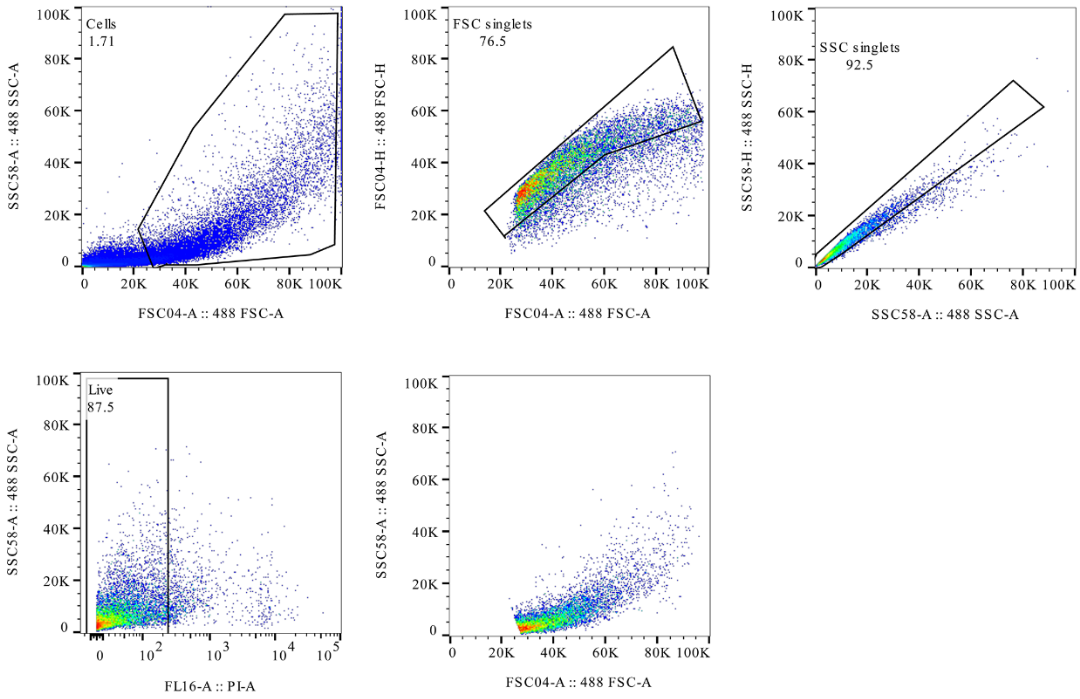

**Supplemental Figure 1. Example of the FACS gating.** (A) FACs gating for total kidney cells. (B) FACS gating for distal convoluted tubule (DCT) cells. For both samples a crude gate is first made to exclude debris and aggregates. Then doublets are removed in first the forward scatter dimension, followed by the side scatter dimension. For total kidney cells (A) the sort gate was on live cells, whereas for DCT cells, the sort gate was on live and GFP positive cells.

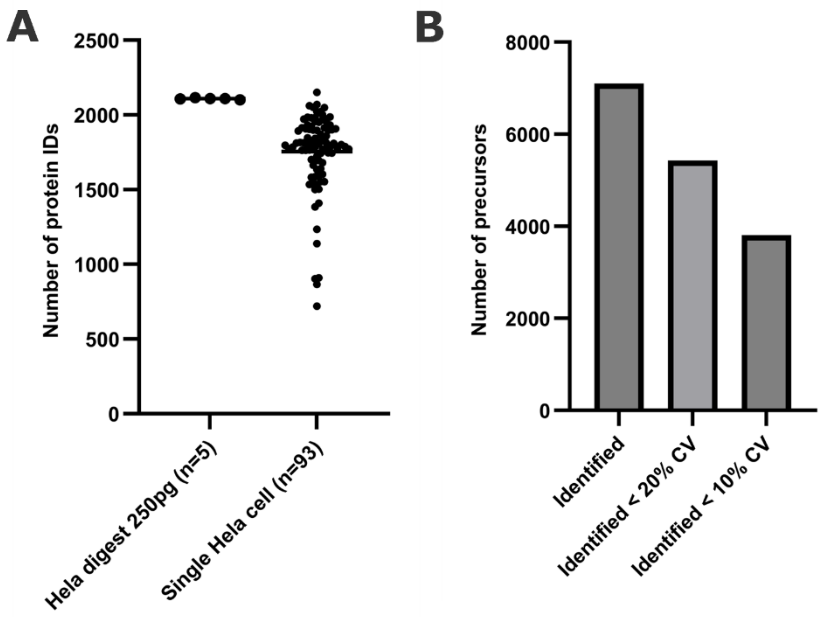

**Supplemental Figure 2. Benchmarking of the LC-MS system and the sample preparation workflow**. (A) Number of identified proteins from 250 pg of Hela digest (n=5) or from single Hela cells (n=93) that went through the entire sample preparation workflow. (B) The number of total identified precursors and those that fulfilled the CV<20% and CV<10% from the 5 runs of 250 pg of Hela digest.

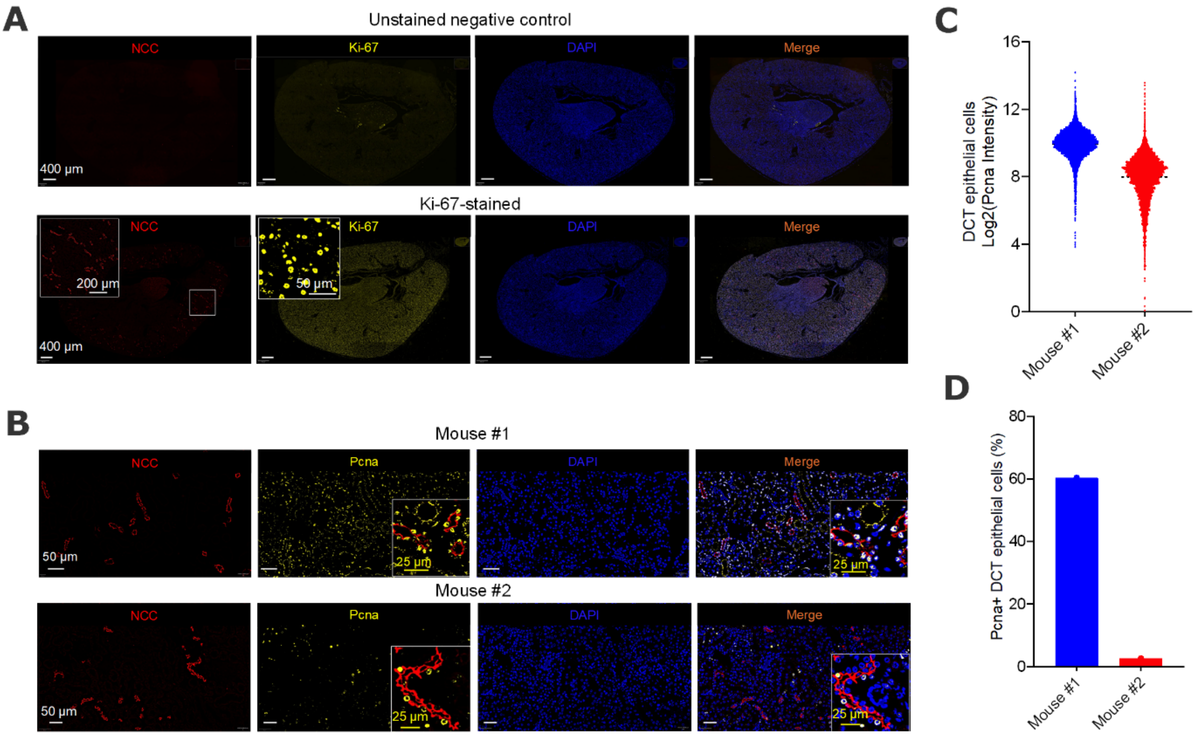

**Supplemental Figure 3. Immunofluorescence analysis of Pcna expression in distal convoluted tubule**. (A) negative control and Ki-67 stained (Figure 4); (B) Representative images of immunofluorescence analyses of NCC and Pcna staining with DAPI counterstain from two mouse kidneys; (C) staining intensities for Pcna within the NCC positive population (DCT) in two mouse kidneys; (D) percentages of Pcna positive DCT cells compared to all DCT cells in two mouse kidneys.
